# Conserved nicotine-activated neuroprotective pathways involve mitochondrial stress

**DOI:** 10.1101/2020.05.28.120576

**Authors:** J. Brucker Nourse, Gilad Harshefi, Adi Marom, Abdelrahaman Karmi, Kim A. Caldwell, Guy A. Caldwell, Millet Treinin

**Affiliations:** Department of Biological Sciences, The University of Alabama, Tuscaloosa, AL, USA; Departments of Neurology and Neurobiology, Center for Neurodegeneration and Experimental Therapeutics, Nathan Shock Center of Excellence in the Basic Biology of Aging, University of Alabama at Birmingham School of Medicine, Birmingham, AL, USA; Department of Medical Neurobiology, Hebrew University – Hadassah Medical School, Jerusalem, Israel

## Abstract

Tobacco smoking is a risk factor for several human diseases. Conversely, smoking also reduces the prevalence of Parkinson’s disease (PD), whose hallmark is degeneration of *substantia nigra* dopaminergic neurons (DNs). We use *C. elegans* as a model to investigate whether tobacco-derived nicotine activates nicotinic acetylcholine receptors (nAChRs) to selectively protect DNs. Using this model we demonstrate conserved functions of DN-expressed nAChRs. We find that DOP-2, a D3-receptor homolog, MCU-1, a mitochondrial calcium uniporter, and PINK-1, PTEN-induced kinase 1, are required for nicotine-mediated protection of DNs. Together, our results support involvement of calcium-dependent mitochondrial stress activation of PINK-1 in nicotine-dependent neuroprotection. This suggests that nicotine’s selective protection of *substantia nigra* DNs is due to the confluence of two factors: first, their unique vulnerability to mitochondrial stress, which is mitigated by increased mitochondrial quality control due to PINK1 activation; and second, to their specific expression of D3 receptors.

## Introduction

Epidemiological studies show that tobacco smoking reduces the prevalence of Parkinson’s disease (PD) (Li et al., 2015). A hallmark of PD is degeneration of *substantia nigra* dopaminergic neurons (DNs). Effects of tobacco smoking on degeneration of these neurons may depend on tobacco-derived nicotine and on nicotinic acetylcholine receptors (nAChRs). Support for this hypothesis comes from several lines of evidence. These include: i) expression studies showing that *substantia nigra* DNs express multiple nAChR subtypes having high affinity for nicotine and thus likely to be affected by the low nicotine concentrations present in the blood of tobacco smokers (Jones, Bolam, & Wonnacott, 2001; Zoli et al., 2002); ii) analysis of PD models showing protective effects of nicotine on *subtantia nigra* DNs (Huang, Parameswaran, Bordia, Michael McIntosh, & Quik, 2009; Y. Liu et al., 2012; Quik, O’Neill, & Perez, 2007); iii) genetic studies implicating RIC3, a chaperone of nAChRs, in PD (Sudhaman et al., 2016); and iv) a recent study that identified *CHRNA6* [encoding a neuronal nAChR subunit selectively affecting DN activity (Quik, Perez, & Grady, 2011)] as a genetic modifier of infantile Parkinsonism (Martinelli et al., 2020).

Several nAChR-dependent processes may explain the neuroprotective effects of nicotine. Nicotine acting through nAChRs was shown to activate protective signaling pathways within neurons (Dineley et al., 2001; Stevens, Krueger, Fitzsimonds, & Picciotto, 2003; Toborek et al., 2007; Yu, Mechawar, Krantic, & Quirion, 2011). Nicotine-mediated neuroprotection may also be a byproduct of altered neuronal activity following chronic nicotine exposure, as reported in smokers (Dani, 2015). Additionally, nicotine may protect neurons indirectly through established neuroinflammation-reducing effects resulting from activation of astrocyte-expressed α7 nAChRs (Jurado-Coronel et al., 2016). But despite such neuroprotective effects, smoking is in fact a known risk factor for cancer, stroke, heart disease, and other prevalent human disorders, including neurodegenerative diseases such as Multiple Sclerosis and Alzheimer’s disease. Mechanisms suggested to reduce prevalence of PD afforded by tobacco smoking should, therefore, also provide plausible explanations for selectivity of these effects (Chang, Ho, Wong, Gentleman, & Ng, 2014; Jiang et al., 2020; Li, Li, Liu, Shen, & Tang, 2015; Otuyama et al., 2019; Rosso & Chitnis, 2019). To date, mechanisms conferring the selective protection to *substantia nigra* DNs, afforded by tobacco smoking, remain undefined.

The nematode *Caenorhabditis elegans* provides a simple and well characterized model for the study of nicotine-mediated protection of DNs. The well described nervous system of *C. elegans* includes eight well characterized DNs (Sawin, Ranganathan, & Horvitz, 2000), which provide an advantageous model for analysis of PD-modifying mechanisms. Exposure to 6-OHDA or MPTP (toxins specifically affecting DNs) or overexpression of human α-synuclein (α-syn; a cause of PD) have similar degeneration-causing effects on *C. elegans* and human DNs (Lakso et al., 2003; Nass et al., 2005; Nass, Hall, Miller, & Blakely, 2002). Research using this model has led to identification and characterization of conserved genes, pathways, and small molecules that function to modulate neurodegeneration (24).

Here, we have undertaken a systematic functional analysis of nAChRs in *C. elegans* DNs. We show that nAChRs alter a dopamine-dependent behavioral response, basal slowing response (BSR), via DN-expressed nAChRs. We also describe nicotine-dependent protection against 6-OHDA toxicity by DN-expressed nAChRs. These results establish *C. elegans* as a conserved model to study the functional effects of nAChRs on DNs, in vivo. Furthermore, we demonstrate the mechanistic utility of this nematode model by identifying nicotine-activated neuroprotective signaling pathways functioning in DNs. Results of this analysis suggest involvement of the mitochondrial stress response, mitochondrial calcium accumulation, and PINK-1 [mutations in which are a cause of PD] in nicotine-mediated protection of DNs (Mouton-Liger, Jacoupy, Corvol, & Corti, 2017). The extensive axons, dense synapses, and high basal activity of *substantia nigra* DNs were suggested to increase metabolic load on these neurons, leading to their increased vulnerability to perturbations in mitochondrial function (Ge, Dawson, & Dawson, 2020). Thus, our findings showing nicotine-induced activation of the PINK-1 mitochondrial quality control pathway may explain selective protection afforded by tobacco smoking to these neurons. Moreover, we show that *C. elegans* DOP-2, a D3-receptor (D3R) homolog, is required for nicotine-mediated protection. D3Rs are selectively expressed in nigrostriatal DNs where they function as auto-receptors (Joyce & Millan, 2007; Yang, Perlmutter, Benzinger, Morris, & Xu, 2020). The involvement of D3Rs in nicotine-mediated protection may further explain the selective protection afforded to DNs by tobacco smoking, as a source of chronic nicotine exposure.

## Results

### RIC-3 and nAChRs function within DNs of *C. elegans* to enhance dopaminergic signaling

*C. elegans* is a simple, genetic model with a well characterized nervous system that can be used to study various cellular phenomena, such as functions of nAChRs in DNs. To determine if nAChRs possess a conserved function in *C. elegans* DNs, as shown for mammalian nAChRs, we examined their effects on the basal slowing response (BSR; reduced locomotion on food), a behavior dependent on dopamine release from DNs (Sawin et al., 2000). RIC-3, a chaperone of nAChRs, is required for maturation of all known nAChRs in *C. elegans* (Halevi et al., 2002; Jospin et al., 2009). Thus, a loss-of-function mutation in *ric-3 (ric-3(lf)* can be used as proxy for effects of loss-of-nAChR function on DN-dependent behaviors. BSR analysis of *ric-3(lf)* animals shows a significant reduction in basal slowing (Figure 1A, B). Rescue experiments in which *ric-3* was expressed selectively in DNs restored basal slowing for *ric-3(lf)* animals (Figure 1C, D). Both *ric-3(lf)* and DN-specific expression of *ric-3* specifically affect locomotion on food but not off food, a result consistent with RIC-3 being required within DNs to increase dopaminergic signaling in response to the presence of food. These results suggest that nAChRs of *C. elegans*, like mammalian nAChRs, affect DN function.

**Figure 1.**
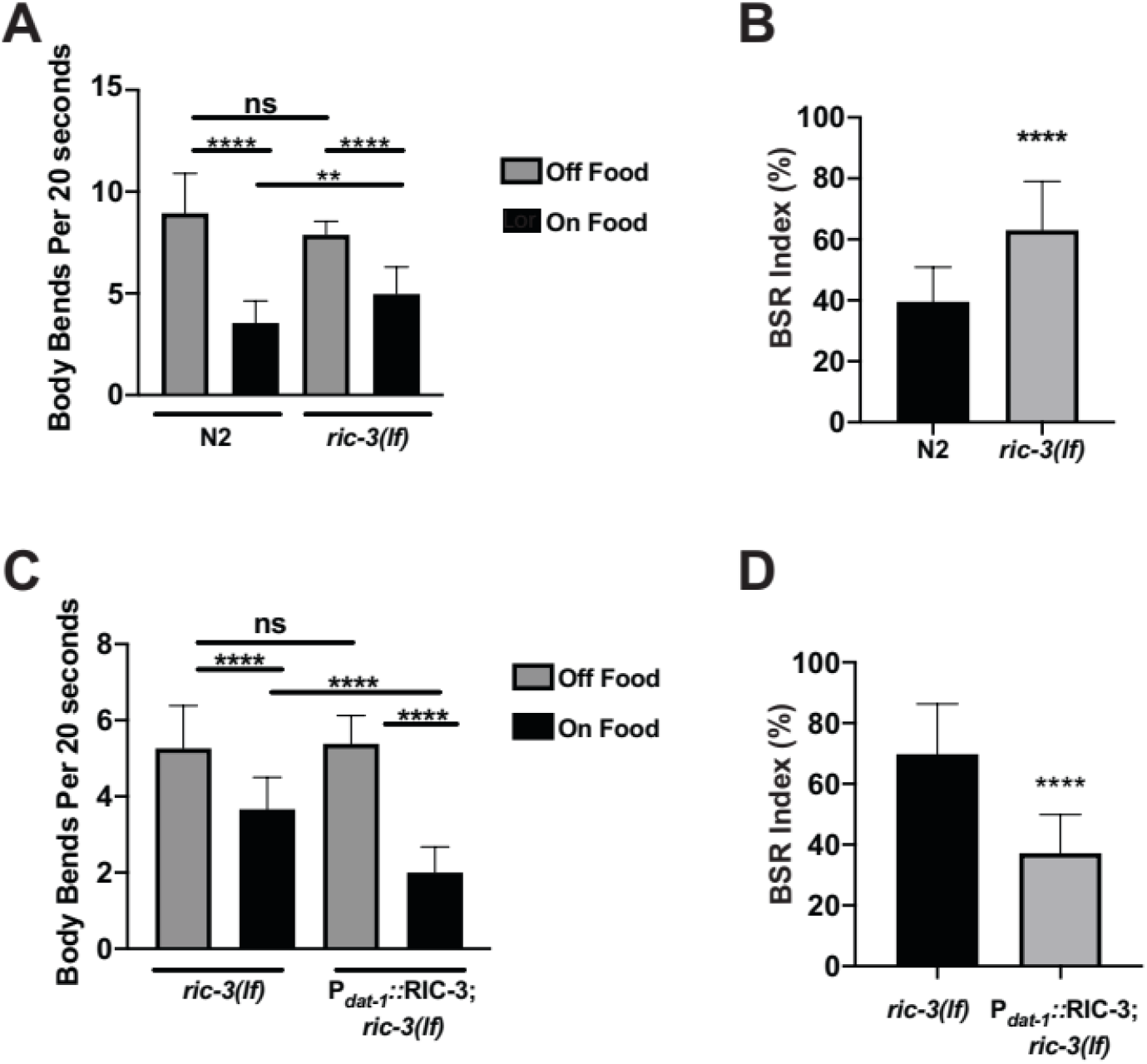
RIC-3 functions in DNs to enhance basal slowing response. (A, B) *ric-3(lf)* exhibit reduced BSR when compared to wild-type (N2) animals. N=2, n=20 animals on and off food. (C, D) Selective expression of RIC-3 in DNs enhances BSR when compared to *ric-3(lf)* animals. N=2, n=13-15 animals on and off food. (A, C) Locomotion speed was measured by number of body bends per 20 seconds on food (black bars) or off food (grey bars). (B, D) BSR index (% body bends per 20 seconds on food per animal relative to average body bends per 20 seconds off food in the same strain and experiment) for data in A or C. Data presented as average + SD; significance was examined using, one-way ANOVA with Tukey’s post hoc multiple comparisons (A, C), or two-tailed unpaired Student’s *t*-test (B, D)); ns - p >0.05, ** - p =0.0063, **** - p <0.0001.

In *C. elegans*, RIC-3 functions to enhance maturation of nAChRs, but not of other ligand-gated ion-channels that were examined (Halevi et al., 2002). Thus, effects of RIC-3 on dopaminergic signaling seen in Figure 1 are likely to be mediated by nAChRs expressed in DNs that require RIC-3 for their functional expression. Furthermore, transcriptomic analysis suggests that seventeen nAChR subunits are expressed in *C. elegans* DNs (J. Cao et al., 2017; Spencer et al., 2011). To examine whether the effects of RIC-3 on dopaminergic signaling is dependent on nAChR activity in DNs, we assayed loss-of-function mutants for various nAChR subunits using the BSR assay (Table 1). Results of this analysis revealed that four of thirteen subunits examined (ligand binding α subunits, UNC-38 and UNC-63, and non-α subunits ACR-2 and LEV-1) are required for proper BSR (Table 1). Figure 2A describes the representative results for the *lev-1(lf)* mutation, wherein BSR was reduced (Figure 2A). These results indicate defective dopamine signaling as quantified by the BSR assay.

**Figure 2.**
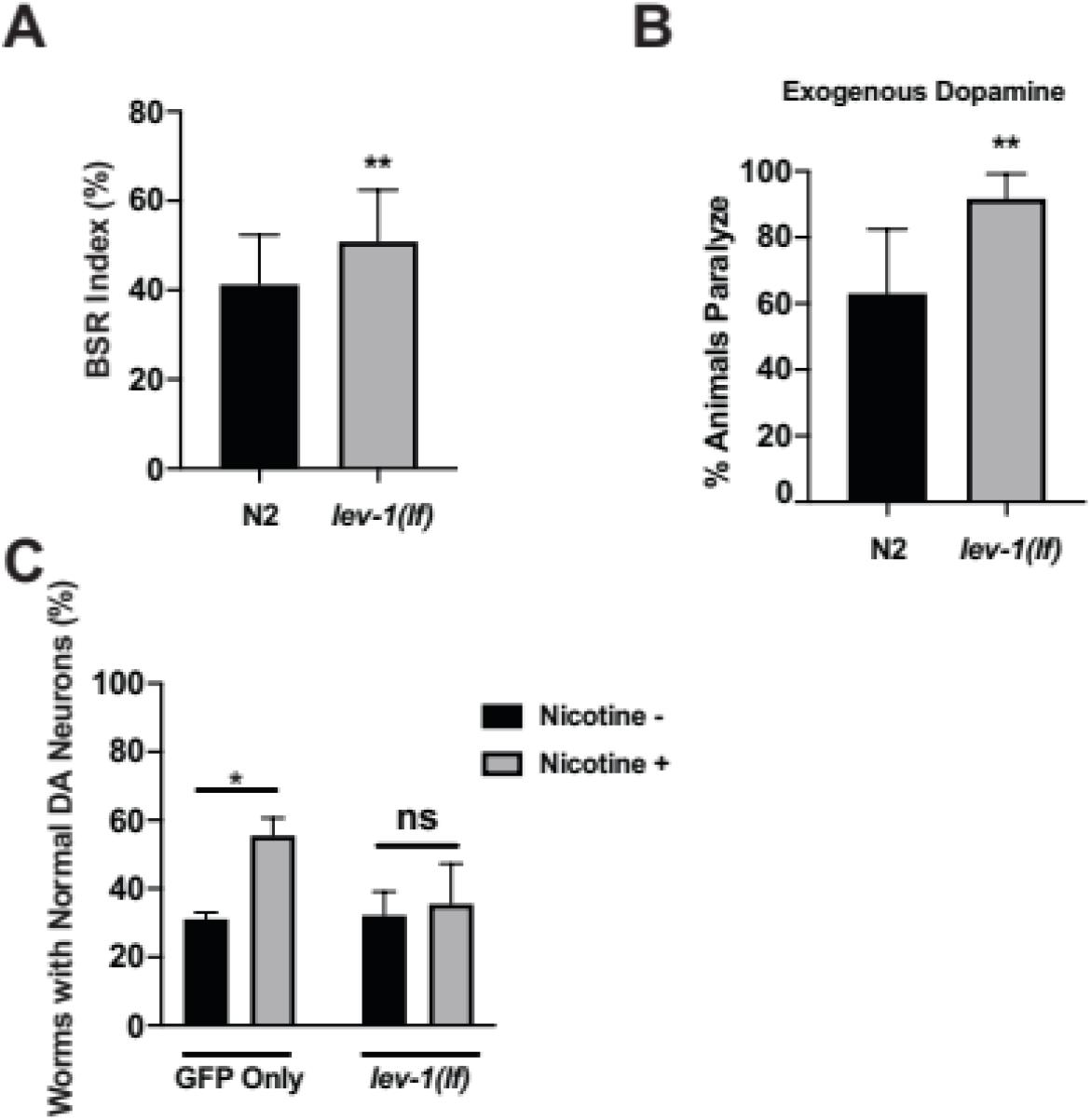
Functions of the LEV-1 nAChR subunit in DNs. (A) *lev-1(lf)* caused basal slowing deficits when compared wild-type animals as shown by increased BSR index (% body bends per 20 seconds on food per animal relative to average body bends per 20 seconds off food in the same experiment for wild-type, N, (N=2, n=30 each) or *lev-1(lf)* (N=2, n=15 each) animals). (B) *lev-1(lf)* animals had increased sensitivity to the paralyzing effects of exogenous dopamine. Results depicted as % animals paralyzed after 20 minutes in the presence of exogenous dopamine, 6 plates, 10 animals per plate. (C) *lev-1(lf)* abolishes nicotine-induced protection against 6-OHDA toxicity. Effects of chronic nicotine exposure on 6-OHDA-induced DN degeneration in *lev-1(lf);* P_*dat-1*_::GFP (*vtIs7*) animals expressing GFP in their DNs is compared to effects of this treatment on the control BY250 strain [P_*dat-1*_::GFP (*vtIs7*)] animals expressing GFP only in DNs N=3, n=30 each. Data represented as average + SD; significance was examined using two-tailed unpaired Student’s *t*-test (A, B) or one-way ANOVA with Tukey’s post hoc multiple comparisons (C); ns - p>0.05, *- p = 0.0149, **- p <0.0097.

**Table 1.** *C. elegans* nAChR subunits examined in this study. Indicated for each subunit: analysis of whether a loss-of-function mutant in the gene displayed reduced BSR (+), no effect (-), or was not determined (n.d.). Additionally, whether the loss-of-function mutants or RNAi knockdown (used for *acr-12, acr-20*) demonstrated a loss-of-nicotine-mediated neuroprotection (+) or not (-) following exposure to 6-OHDA. ^**1**^Expression information for these subunits in DNs was compiled from reports of Spencer et al., 2011 (+) and Cao et al., 2017 (*). It should be noted that occasional discrepancies between these two expression studies exist and might have resulted from the use of different developmentally staged animals or from the profiling method’s sensitivity and/or signal-to-noise ratio.

| **nAChR subunit** | **BSR reduced** | **Nicotine-mediated neuroprotection** | **Expressed in DNs<sup>1</sup>** |
| --- | --- | --- | --- |
| *unc-38* | + | - | +/* |
| *unc-63* | + | - | +/* |
| *acr-2* | + | - | +/- |
| *lev-1* | + | + | +/* |
| *acr-12* | - | + | +/* |
| *acr-20* | n.d. | + | +/* |
| *unc-29* | - | + | +/- |
| *acr-5* | - | n.d. | +/* |
| *acr-8* | - | n.d. | +/- |
| *acr-15* | - | n.d. | +/- |
| *acr-18* | - | n.d. | +/- |
| *acr-21* | - | n.d. | +/- |
| *deg-3* | - | n.d. | +/* |
| *des-2* | - | n.d. | +/* |

All four subunit mutants affecting BSR were also examined for their sensitivity to exogenous dopamine. In *C. elegans*, the application of exogenous dopamine causes paralysis of wild-type animals (Chase, Pepper, & Koelle, 2004). None of the mutations examined reduced the paralyzing effects of exogenous dopamine. In fact, *lev-1(lf)* even showed enhanced responsiveness to exogenous dopamine (Figure 2B). Considering these data, reduced BSR in these mutants is likely to be a result of decreased dopamine release from DNs and not of reduced responsiveness to dopamine. This together with the results on RIC-3 functioning in DNs to affect BSR, strongly support a conserved role for nAChRs in modulating DN-activity and dopamine release from these neurons.

### Nicotine activation of nAChRs protects *C. elegans* DNs from 6-OHDA toxicity

Nicotine was shown to protect mammalian DNs from 6-OHDA, a DN-specific neurotoxin, in a nAChR dependent manner (Ryan, Ross, Drago, & Loiacono, 2001). In order to determine if the neuroprotective effects of nicotine on DNs are conserved in *C. elegans*, we treated animals chronically for 96 hours with nicotine and examined effects of this treatment on 6-OHDA-induced neurodegeneration (Figure 3A). The nicotine concentration selected (62μM) was previously shown to significantly affect gene expression but not fertility nor viability of *C. elegans* (Polli et al., 2015; Smith et al., 2013). Our results demonstrate that animals treated only with nicotine have no change in DN morphology (Figure 3B, D), signifying nicotine has no deleterious effect at the tested concentration. 6-OHDA only treatments (Figure 3B, E) caused significant neurodegeneration, while chronic exposure to nicotine provided robust neuroprotection from 6-OHDA (Figure 3B, F). These data demonstrate that nicotine significantly protects DNs from the selective neurodegeneration induced by 6-OHDA.

**Figure 3.**
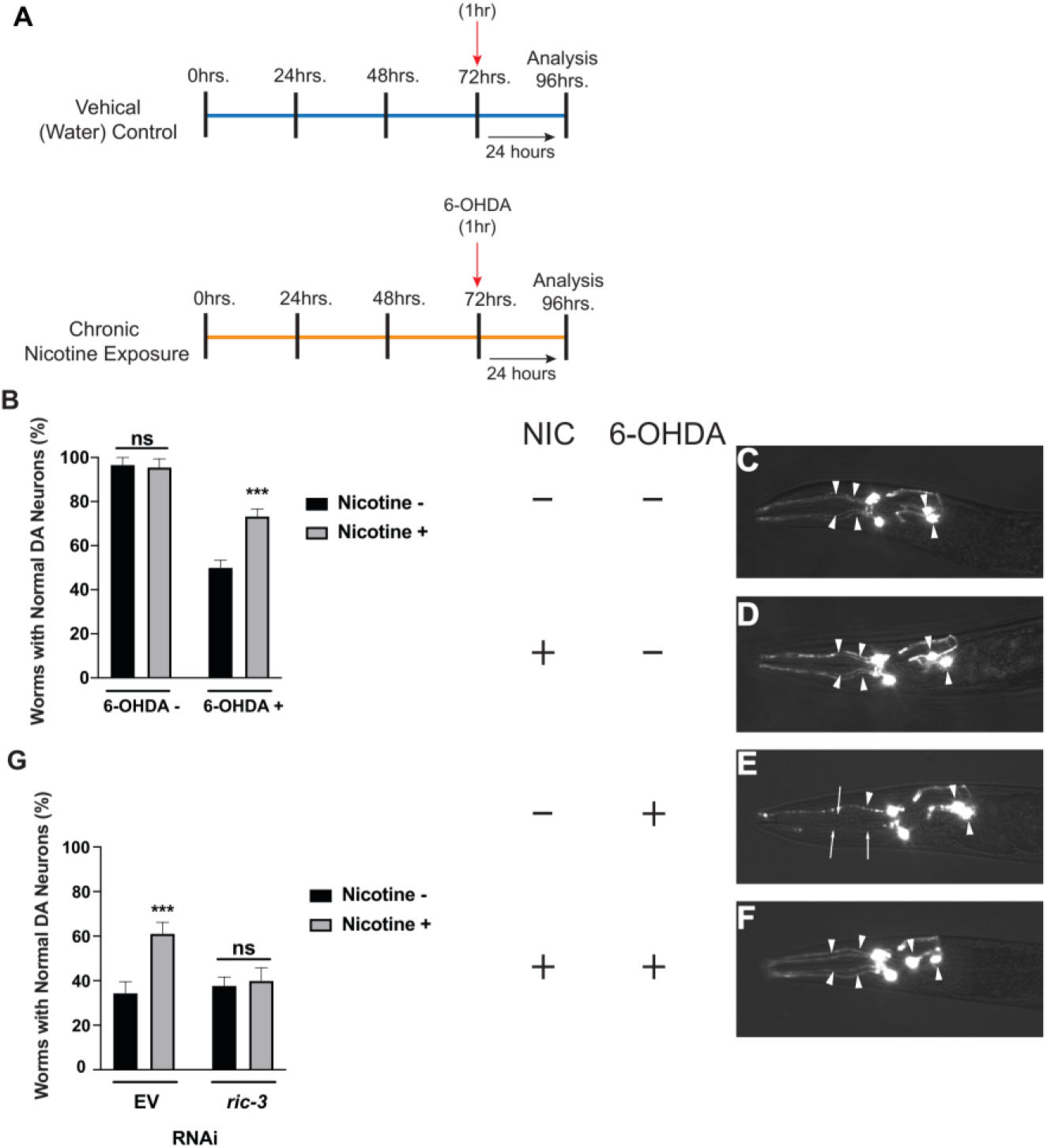
Chronic nicotine treatment protects DNs in a RIC-3 dependent manner. (A) Timeline of experiments. (A-Top) Control experiments without nicotine exposure. (A-Bottom) Animals in nicotine group exposed to 62μM nicotine from hatching, during and after 6-OHDA exposure. Degeneration of DNs was scored twenty-four hours after 10mM 6-OHDA/2mM ascorbic acid exposure or 2mM ascorbic acid alone. (B) Chronic nicotine treatment attenuates 6-OHDA-induced DN degeneration in BY250 worms expressing only GFP in their DNs [P_*dat-1::*_GFP (*vtIs7*)]. (C-F) Representative images matching the conditions of panel B. (C) Solvent (ddH_2_O/2mM ascorbic acid) control with all six anterior DNs intact (arrowheads). (D) Nicotine only treatment with all six anterior DNs intact (arrowheads). (E) 6-OHDA-induced DN degeneration, with three degenerated (arrows) and three intact (arrowheads) DNs depicted. (F) Nicotine treatment of 6-OHDA treated animals with all six anterior DNs intact and protected from 6-OHDA toxicity (arrowheads). (G) DN-specific RNAi knockdown of *ric-3* (using UA202 worms) abolishes nicotine-induced protection against 6-OHDA toxicity compared to empty vector (EV) control. Data is represented as average + SD (N=3, n= 30 each). Significance was examined using two-way ANOVA with Tukey’s post hoc multiple comparison test (A, F); ns - p >0.05, *** - p <0.0008.

We explored additional nicotine exposure paradigms, including a one-hour acute exposure that was co-administered with 6-OHDA, and a one-hour pre-exposure to nicotine, which was administered 24 hours before the 6-OHDA treatment and that was combined with acute exposure (Figure 4A). As shown in Figure 4 (B,C), acute nicotine treatment did not provide significant protection to DNs whereas adding pre-treatment to acute exposure was protective, albeit at a significance level lower than that observed with chronic nicotine exposure (p = 0.0178 vs. p = 0.0002, respectively) (Figure 4C vs. 3B). Therefore, all subsequent experiments utilized chronic treatment with nicotine to maximize its protective effects. Notably, these two paradigms of nicotine-induced neuroprotection in *C. elegans* (chronic and pre-exposure) have also been observed to be neuroprotective in mammalian experimental models (Huang et al., 2009; Janson, Fuxe, & Goldstein, 1992). The long lasting (24 hours) effects of nicotine pre-exposure is consistent with nicotine-dependent gene expression changes enabling or enhancing nicotine-induced protection of DNs.

**Figure 4.**
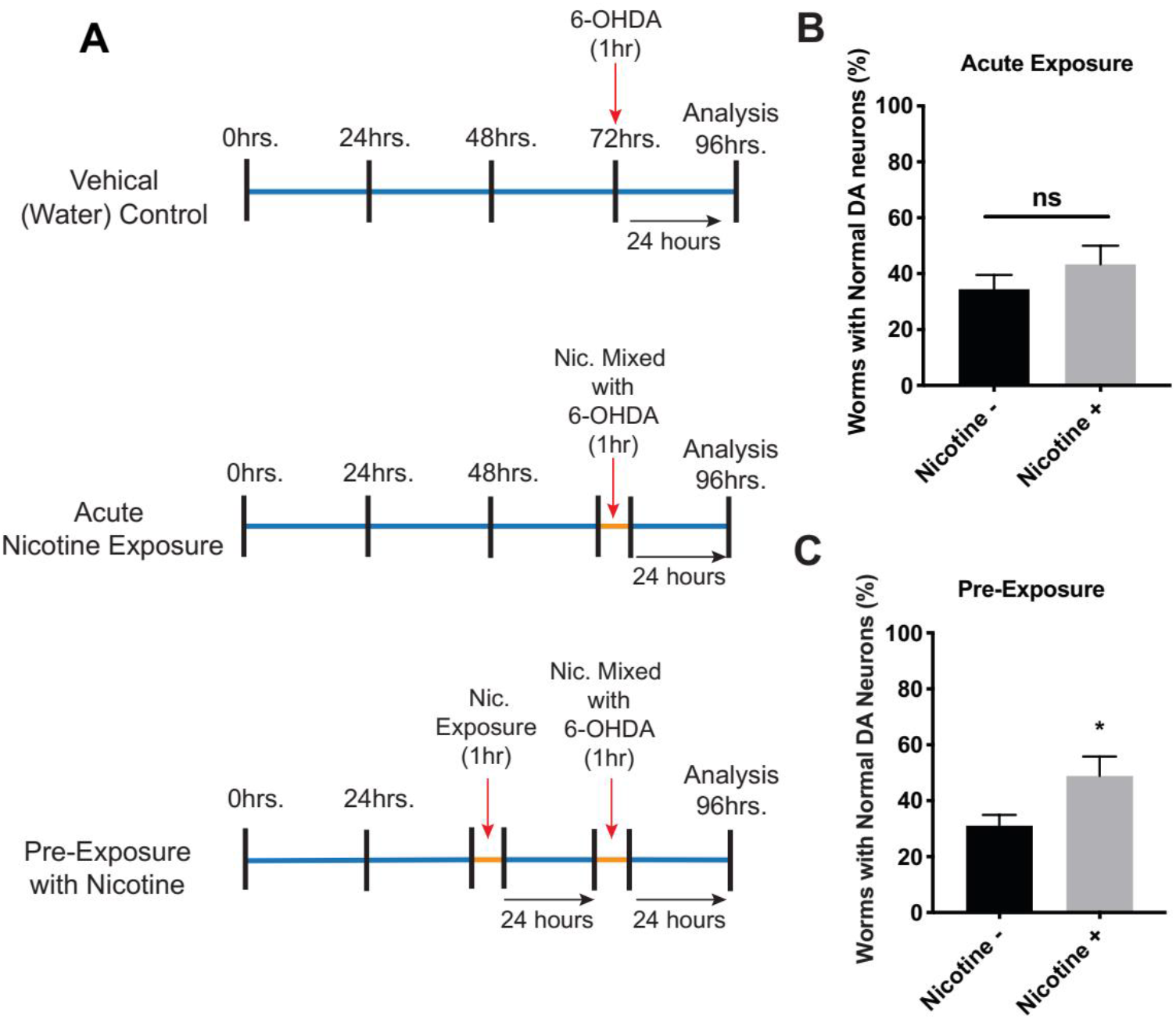
Pre-exposure to nicotine protects DNs from 6-OHDA. (A) Timeline depicting nicotine treatments protocols. (A-Top) Control protocol. (A-Middle) Acute exposure, addition of 62μM to the assay mix only during 6-OHDA treatment. (A-Bottom) Pre-exposure, one-hour pre-exposure to 62μM nicotine the day before 6-OHDA + nicotine (acute exposure). Examined in BY250 worms expressing GFP only in DNs. (B) Acute nicotine is not neuroprotective. (C) A nicotine pre-exposure + acute exposure is protective against 6-OHDA toxicity. Data represented as average + SD (N=3, n=30 worms each). Significance was examined using two-tailed unpaired Student’s *t*-test; ns - p > 0.05, * - p= 0.0178.

To determine if the expression of nAChRs in DNs is required for nicotine-induced neuroprotection in *C. elegans*, we knocked down *ric-3* selectively in DNs. RNAi silencing in DNs was accomplished by the RNAi feeding method using a *sid-1(lf)* mutant strain that only expresses SID-1, a dsRNA transporter, in DNs (Harrington, Yacoubian, Slone, Caldwell, & Caldwell, 2012). As depicted in Figure 3G, DN-specific knockdown of *ric-3* in animals not treated with nicotine does not affect 6-OHDA-induced neurodegeneration when compared to the empty vector (EV) RNAi control. However, *ric-3* silencing does abolish the neuroprotective effects of nicotine on animals exposed to 6-OHDA when compared to the nicotine-treated EV control (Figure 3G). Silencing of *ric-3* in DNs not treated with 6-OHDA did not impact DN morphology (not shown). Elimination of nicotine-induced neuroprotection following selective *ric-3* silencing in DNs implies a vital role for DN-expressed nAChRs to convey the protective effects of nicotine.

To further investigate whether nAChRs are required for protective effects of nicotine on DNs against 6-OHDA toxicity, we individually tested seven nAChR subunits. These subunits were examined by either DN-specific RNAi knockdown or loss-of-function mutant alleles. The seven subunits examined included the four subunits shown to be needed for proper BSR (Table 1) – subunits likely to function in DNs to affect dopamine release – and three additional subunits shown to be expressed in the DNs through transcriptomics data (J. Cao et al., 2017; Spencer et al., 2011). This approach identified four subunits (α subunits ACR-12 and ACR-20, and non-α subunits LEV-1 and UNC-29) that are necessary for nicotine-induced neuroprotection (Table 1). Notably, *lev-1(lf)* was the only mutant found to impact both dopamine signaling (Figure 2A), and nicotine-induced DN protection (Figure 2C). Together, these results demonstrate that, similar to mammals, nicotine elicits a neuroprotective effect on *C. elegans* that is dependent on nAChR expression in DNs.

Collectively, our results suggest that *C. elegans* DNs express at least two distinct nAChRs; 1) one affecting DN activity and dopaminergic signaling, and 2) the other mediating the protective effects of nicotine. Evidence for this hypothesis comes from the following considerations: nAChRs are pentamers (heteromers or homomers), but we identified seven subunits involved in either dopamine signaling or neuroprotection. Moreover, only one subunit, LEV-1, was shown to affect both DN activity and DN protection; ACR-2, UNC-38 and UNC-63 only affect BSR while ACR-12 and UNC-29 only affect nicotine-mediated protection of DNs. No viable mutant is available for *acr-20*, thus, this subunit was only examined for its effects on nicotine-induced protection of DNs, using DN-selective RNAi silencing. Interestingly, this subunit was shown to function as a homomeric receptor and thus may form a third DN-expressed nAChR (Baur, Beech, Sigel, & Rufener, 2015).

### Conserved mechanisms enable the protective effects of nicotine

Studies involving mammalian cell cultures have identified several genes and pathways mediating nicotine-induced neuroprotection. One such study showed that the neuroprotective effects of nicotine involve increased cytosolic calcium, voltage-gated calcium channels, and the calcium and calmodulin dependent phosphatase, calcineurin (Stevens et al., 2003). To determine whether these mechanisms are conserved, we examined effects of nicotine-induced neuroprotection following DN-specific knockdown of *egl-19*, encoding a voltage-gated calcium channel, and of *tax-6*, encoding the *C. elegans* homolog of calcineurin. These experiments were performed using a strain enabling selective RNAi in DNs (Harrington et al., 2012). Animals were raised on RNAi bacteria for two generations (P0 and F1), with only the F1 generation receiving chronic treatment of nicotine (Figure 5A). In this, and the following RNAi experiments, *ric-3* RNAi was used as an internal control. Previous studies demonstrated that the individual knockdown of *egl-19* or *tax-6* caused DN-protection from neurodegeneration induced by α-synuclein, a protein implicated in human PD, thus implying degeneration enhancing activities of these two genes in DNs (Caraveo et al., 2014). Similarly, when *egl-19* or *tax-6* are knocked-down selectively in DNs, we show reduced 6-OHDA-dependent neurodegeneration, even without nicotine treatment (Figure 5B, C), thus supporting a role for these two genes in enhancing degeneration of DNs. This protective effect is not additive when combined with nicotine, suggesting that voltage-gate calcium channels (EGL-19; Figure 5B) and calcineurin (TAX-6; Figure 5C) function downstream of nAChR activation. However, lack of additivity is not sufficient evidence for this pathway as we cannot rule out a ceiling effect masking additivity.

**Figure 5.**
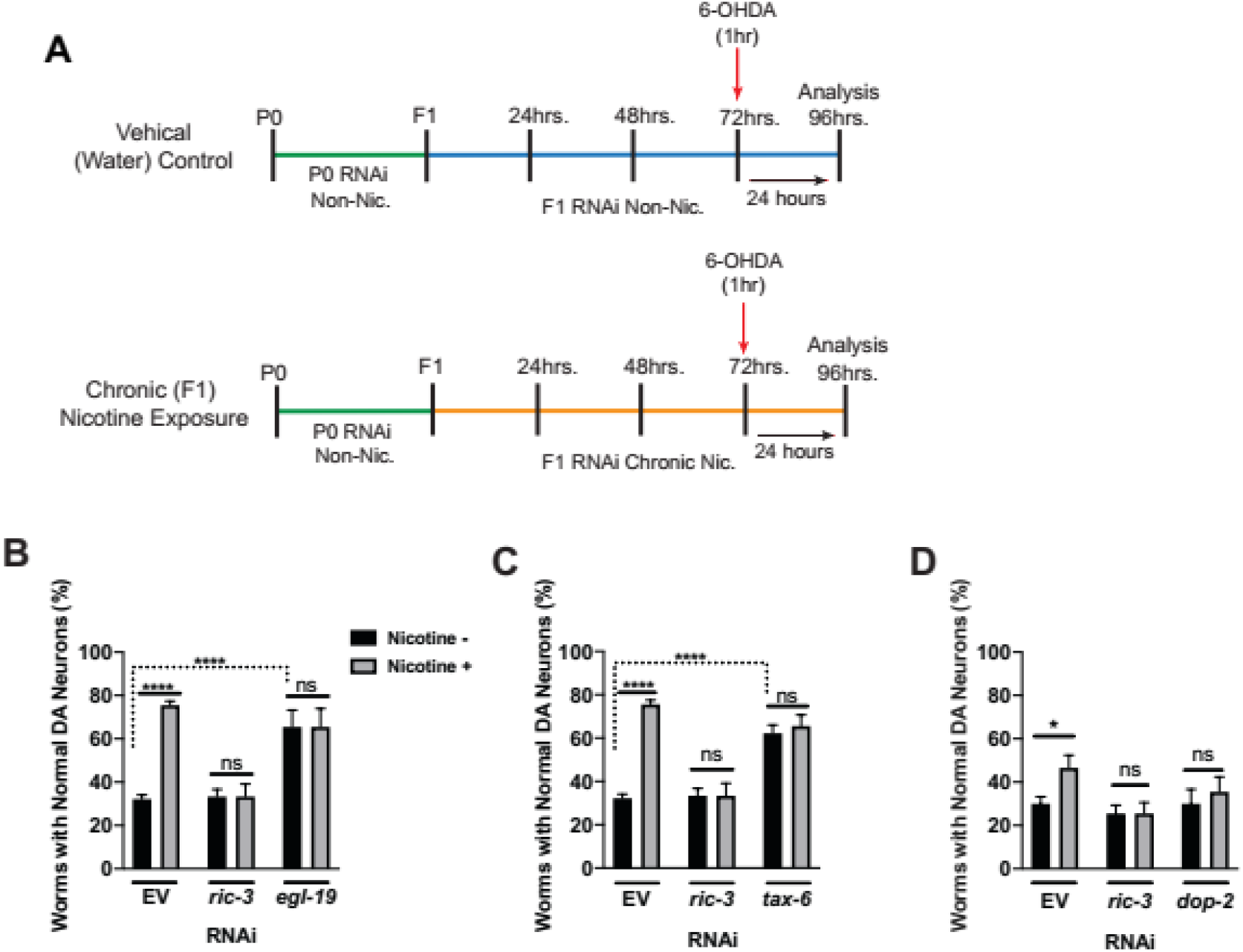
Conserved mechanisms mediate protective effects of nicotine. (A) Timeline depicting RNAi and chronic nicotine treatments. Animals were raised on RNAi bacteria for two generations (P0 and F1). Only the F1 generation were exposed to nicotine. (B-D) RNAi experiments involving the knockdown of *ric-3* or *egl-19* (B) or *tax-6* (C) or *dop-2* (D) in the DN-specific RNAi-sensitive strain, UA202, when animals were grown in the presence (grey bars) or absence (black bars) of 62μM nicotine and exposed to 10mM 6-OHDA/2mM ascorbic acid. (B) *egl-19* and (C) *tax-6* knockdown alone protected DNs from 6-OHDA toxicity and nicotine had no additional effects. (D) *dop-2* knockdown abolished nicotine-induced protection from 6-OHDA. Data shown as average + SD (N=3, n= 30 animals each). Significance was examined using two-way ANOVA with Tukey’s post hoc multiple comparison test; ns - p >0.05, **** - p <0.0001, * - p = 0.0259.

Another signaling protein shown to enable nicotine-induced protection of mammalian DNs is the dopamine D3-receptor (D3R) (Bono et al., 2019); an effect requiring its interaction with the β2 nAChR subunit (Bontempi, Savoia, Bono, Fiorentini, & Missale, 2017). The *C. elegans* dopamine receptor, DOP-2, shares many similarities with D3R including being expressed in DNs, functioning via G_i_/G_o_ to inhibit adenylyl cyclase, and functioning as an auto-receptor to inhibit dopamine release (Formisano et al., 2020; Suo, Sasagawa, & Ishiura, 2003). To examine whether DOP-2 functions as part of the protective signaling pathway activated by nicotine, we knocked down *dop-2* specifically in DNs. We found that, like *ric-3, dop-2* knockdown does not enhance neurodegeneration, but its expression in DNs is required for nicotine-mediated neuroprotection (Figure 5D). This result suggests conservation of the interaction between nAChRs and the D3R/DOP-2 in nicotine-induced protection.

Our results show that, in *C. elegans*, DOP-2 function is required for nicotine-induced protection of DNs, a result consistent with studies involving human D3R signaling in DNs (Bontempi et al., 2017). Our data also support involvement of calcium-dependent signaling in nicotine-induced neuroprotection similarly to what was shown in mammals. Taken together, we demonstrate conservation of mechanisms enabling nicotine-induced neuroprotection between *C. elegans* and mammal (Stevens et al., 2003).

### Nicotine-induced neuroprotection depends on the mitochondrial stress response

By stimulating nAChRs, nicotine was shown to affect mitochondrial function and induce a mitochondrial stress response (Gergalova et al., 2012; Lykhmus et al., 2014; Zahedi et al., 2019). To examine whether chronic nicotine exposure impacts mitochondrial stress in *C. elegans*, we used an established mitochondrial stress reporter, *hsp-6::GFP* (Yoneda et al., 2004). Our results depicted significantly higher *hsp-6::GFP* expression in nicotine-treated animals. These results show systemic increase in the mitochondrial stress response resulting from chronic nicotine exposure on days four and seven post-hatching when compared to worms not exposed to nicotine (Figure 6A).

**Figure 6.**
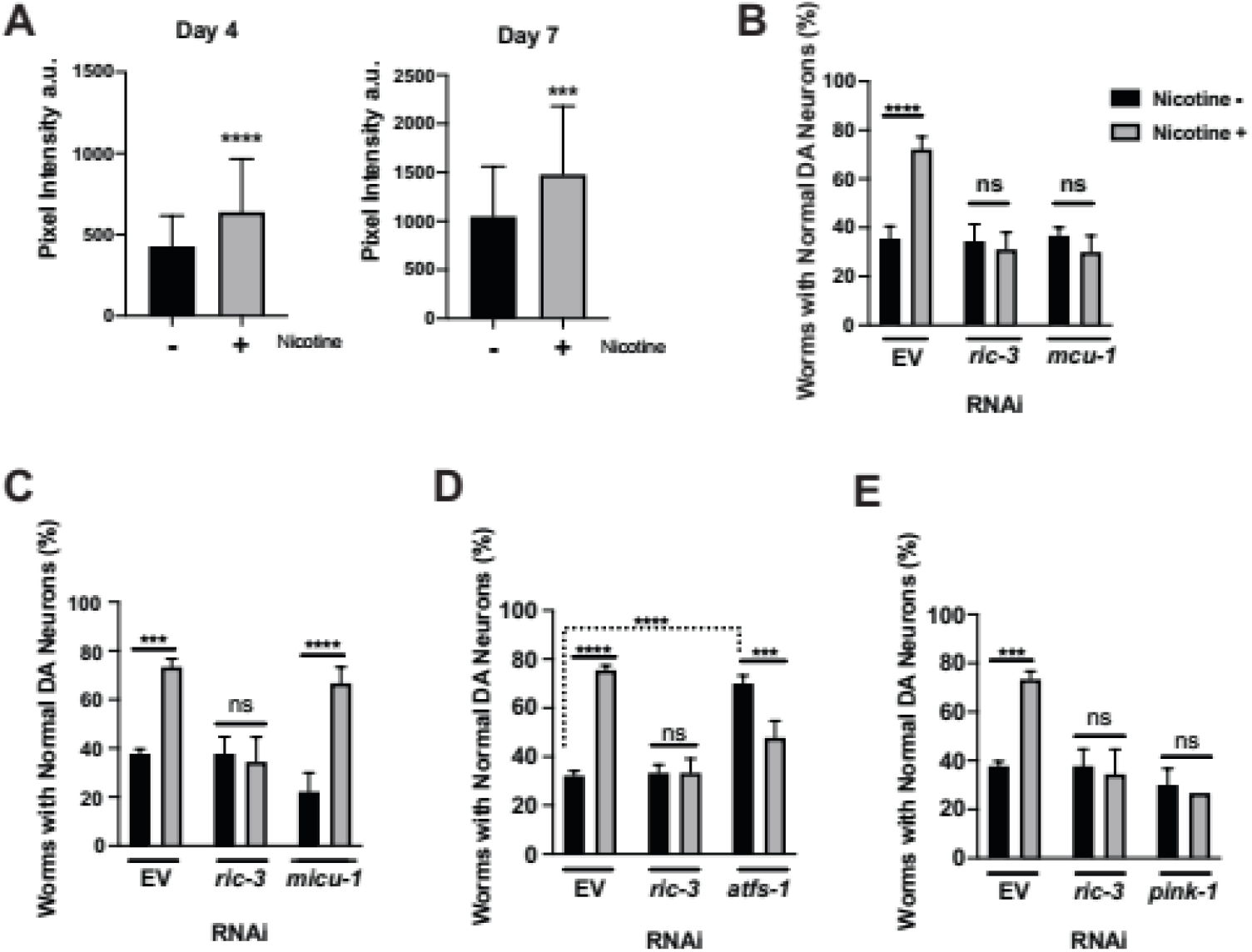
Protective effects of nicotine and mitochondrial stress. (A) A *hsp-6::GFP* reporter strain chronically treated with 62μM nicotine showed increased mitochondrial stress (N=2-3, n=30 each). (B-E) RNAi experiments knocking down gene targets in the DNs of selectively RNAi-sensitive UA202 animals grown in the presence (grey bars) or absence (black bars) of 62μM nicotine and exposed to10mM 6-OHDA/2mM ascorbic acid (N= 3, n=30 each). (B) *mcu-1* knockdown abolished nicotine-induced protection. (C) *micu-1* knockdown had no effect on nicotine-induced protection. (D) *atfs-1* knockdown alone protects against 6-OHDA toxicity, while the addition of nicotine exposure decreased its protective effects. (E) *pink-1* knockdown abolished nicotine-induced protection. Data shown as average + SD and significance was examined using two-tailed t-test (A), or two-way ANOVA with Tukey’s post hoc multiple comparison test (B-E); ns - p >0.05, *** - p <0.0004, **** - p <0.0001.

Our results demonstrate that the relationship between nicotine and mitochondrial stress response is conserved between *C. elegans* and mammals (Figure 6A). This relationship, however, has not been previously linked to nicotine-induced neuroprotection. To investigate whether pathways known to mediate responses to mitochondrial stress also modulate the neuroprotective effects of nicotine, we examined four genes shown to influence the mitochondrial stress response and to function in *C. elegans* DNs, as demonstrated by their effects on α-synuclein-dependent neurodegeneration (*mcu-1, micu-1*, *atfs-1*, and *pink-1*) (Martinez, Kim, Ray, Caldwell, & Caldwell, 2015; Martinez et al., 2017).

MCU-1 is calcium uniporter involved in mitochondrial calcium import and homeostasis. MICU-1 is a calcium sensor responsible for coupling increased cytosolic calcium to increased activity of the MCU-1 (Pendin, Greotti, & Pozzan, 2014). The silencing of *mcu-1* in animals not exposed to nicotine did not enhance neurodegeneration when compared to the EV control, but *mcu-1* silencing neutralized the neuroprotective effects of nicotine against 6-OHDA, when compared to the nicotine-treated EV control (Figure 6B). Knockdown of *micu-1* did not enhance neurodegeneration of animals not treated with nicotine and did not reduce nicotine-mediated neuroprotection (Figure 6C). Thus, while MCU-1 is required for nicotine-induced protection of DNs, MICU-1 is not. This suggests that nicotine induces calcium influx into mitochondria through MCU-1 in a manner independent of MICU-1 function. Interestingly, MICU -/- knockout mice, that exhibit severe mitochondrial defects in calcium homeostasis and developmental abnormalities, show greatly improved mitochondrial calcium parameters as they age, suggestive of MCU1-associated functional remodeling in the absence of MICU1 (J. C. Liu et al., 2016). It is also clear that differential expression of uniporter regulatory proteins, like MICU1, may account for functional tuning of calcium uptake in a tissue-specific manner (Paillard et al., 2017). It is intriguing to speculate that, in a disease of aging like PD, the neuroprotective selectivity of DNs in response to nicotine may reflect a functional change in uniporter dynamics that can both respond to, and influence, mitochondrial stress.

To examine if inducing mitochondrial stress is necessary for the protective effects of nicotine, we examined its dependency on ATFS-1 activity. This protein is a transcription factor that shuttles to the nucleus as part of the mitochondrial unfolded protein response (UPR^mt^) to activate protective gene expression (Nargund, Pellegrino, Fiorese, Baker, & Haynes, 2012). Knockdown of this gene in DNs produced a paradoxical increase in protection against α-synuclein-dependent neurodegeneration, suggesting that the ATFS-1 signaling pathway may have deleterious effects when chronically activated (Martinez et al., 2017). Similarly, *atfs-1* knockdown in animals not treated with nicotine protected DNs from 6-OHDA toxicity (Figure 6D). Interestingly, these effects are not additive to the protective effects of nicotine. Instead, the combined effects of *atfs-1* knockdown and chronic nicotine exposure resulted in a significant decrease, but not a complete loss, of neuroprotection (Figure 6D). This suggests an interaction between two parallel pathways that impinge on a common downstream target process. A likely explanation for the reduced protection seen when activating both pathways is a bell-curve shaped effect, whereby protection is achieved upon limited activation of this downstream process, but both higher and lower activity are detrimental. Such bell-curve shaped effects are also seen for the effects of cytosolic calcium, which at low levels is necessary for normal physiology, but at higher levels is toxic (Choi, 1985). This switch between neuroprotection and neurodegeneration is also a characteristic of calcineurin activity in DNs (Caraveo et al., 2014).

Lastly, we examined the interaction of nicotine with PINK-1, a kinase which is stimulated by mitochondrial stress to activate mitophagy and additional protective processes mostly involving mitochondrial quality control (Ge et al., 2020; Mouton-Liger et al., 2017). RNAi silencing of *pink-1*, without nicotine, does not enhance dopaminergic neurodegeneration when compared to the EV control but it has been shown to do so in presence of other neuronal stressors, such as α-synuclein overexpression in DNs (Martinez et al., 2015). Here, notably, *pink-1* silencing neutralizes the neuroprotective effects of nicotine against 6-OHDA toxicity when compared to the nicotine-treated EV control (Figure 6E). Thus, PINK-1 is likely to function downstream of nAChR activation in DNs to enable the neuroprotective effects of chronic nicotine exposure. This represents a previously unreported mechanism whereby nicotine modulates survival of DNs in vivo via activation of PINK-1, for which loss-of-function is a heritable cause of PD.

## Discussion

Epidemiological studies show that tobacco smoking reduces the prevalence of PD by 40-50% (Ma, Liu, Neumann, & Gao, 2017). This strikingly protective effect of smoking is surprising considering the multiple epidemiological studies indicating smoking to be a risk factor for many diseases, including neurodegenerative diseases such as Multiple Sclerosis (MS) and Alzheimer’s disease (Chang et al., 2014; Jiang et al., 2020; Rosso & Chitnis, 2019). Protective mechanisms suggested to reduce prevalence of PD in smokers should, therefore, explain the selective protection of *substantia nigra* DNs documented among smokers. In addition, such mechanisms should further our understanding of molecular factors that impact the survival of DNs as a selectively vulnerable neuronal subclass in PD. In this study, we identify molecular mechanisms enabling the effects of tobacco-derived nicotine on DNs, mechanisms that may explain selective protection of *substantia nigra* DNs afforded by this activity.

Neuroprotective effects of tobacco-derived nicotine and nAChRs was suggested as an explanation for the reduced prevalence of PD in smokers. Indeed, previous work has shown that nicotine and nAChRs protect neurons from various insults (Picciotto & Zoli, 2008). Here, we demonstrated nicotine-induced protection of *C. elegans* DNs is dependent on DN-expressed nAChRs. Thus, revealing conservation of nicotine-induced neuroprotection between mammals and *C. elegans*. Our results show that *C. elegans* nAChRs are needed both for DN activity, and thus for dopaminergic signaling, as well as for nicotine-induced neuroprotection of DNs. We identified seven nAChR subunits that are needed for either DN activity and/or nicotine-induced neuroprotection against 6-OHDA toxicity. Only one of these subunits affects both nAChR-dependent activities. Thus, our results suggest distinct DN-expressed nAChRs affecting DN activity or DN survival.

Prior studies suggested that interactions between nAChRs and D3Rs are required for nicotine-mediated protection of human DNs (Bono et al., 2019). D3Rs are selectively expressed in DNs and agonists for D3Rs were shown to protect and even restore neuronal functions disrupted in PD (Joyce & Millan, 2007). Here, we show that nicotine-mediated protection following knockdown of *dop-2*, a *C. elegans* D3R homolog (Formisano et al., 2020), is completely abolished. We suggest that nicotine targets protective D3R-dependent signaling pathways to selectively protect DNs (Figure 7). Thus, our results present a plausible explanation for enhanced nicotine-mediated protection that is limited to DNs.

**Figure 7.**
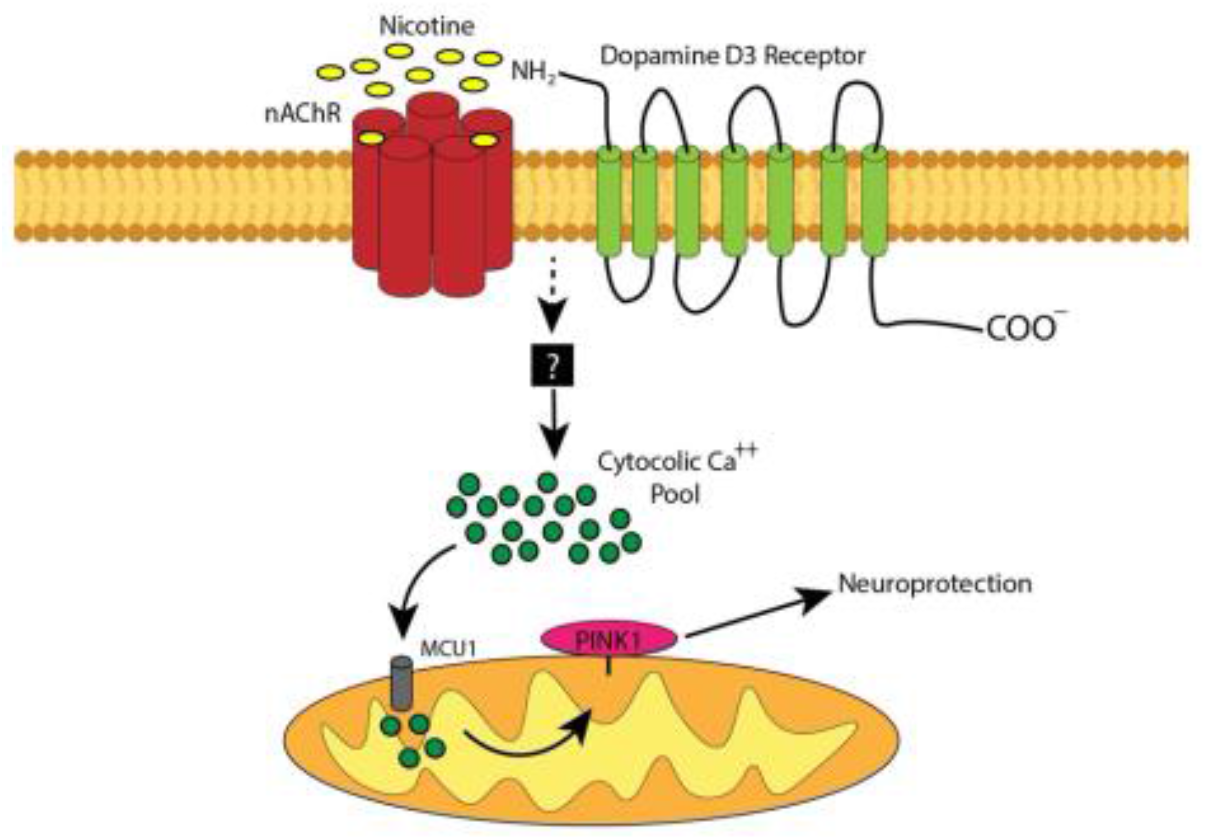
Proposed mechanism enabling nicotine-mediated protection of DNs. Effects of nicotine are mediated by a protective nAChR interacting with the dopamine D3-receptor (D3R) on the plasma membrane to increase cytosolic calcium levels. Cytosolic calcium is sequestered by the mitochondrial calcium uniporter (MCU1), thus resulting in increased mitochondrial calcium levels, mitochondrial stress, and activation of PINK1-associated neuroprotection.

Together, our results establish *C. elegans* DNs as a conserved model enabling investigations of nicotine-induced neuroprotective pathways. Mammalian studies show that nicotine, through nAChR-activation, affects several signaling pathways. These pathways include calcium signaling and the mitochondrial stress response. In the present study, we examined the role of these pathways in nicotine-induced protection of DNs using this newly established model.

Based on our results with *egl-19* and *tax-6* knockdown, we suggest that voltage-gated calcium channels (EGL-19) and the calcium/calmodulin-dependent phosphatase (TAX-6), calcineurin, function downstream of nAChR activation. Studies in mammalian cell cultures suggested that voltage-gated calcium channels are needed at two different stages. First, they function downstream of nAChR-dependent depolarization to amplify calcium signals. Subsequently, they function as a target of calcineurin, which following calcium entry, dephosphorylates and inactivates the channel (Stevens et al., 2003). Our results are consistent with both possibilities. However, only the second explanation is consistent with the large protective effects of *egl-19* knockdown in the absence of nicotine. Future work should examine whether in *C. elegans* calcineurin (TAX-6) dephosphorylates EGL-19 to inhibit its activity, thus, reducing calcium influx into DNs.

Additional analysis supports involvement of the calcium uniporter (MCU-1), enabling calcium influx into mitochondria, in nicotine-mediated protection. We show that nicotine activates mitochondrial stress in *C. elegans* and that the neuroprotective effects of nicotine require PINK-1, a protein known to be activated by mitochondrial membrane depolarization (Mouton-Liger et al., 2017). Our results imply that a nicotine-activated PINK-1-dependent pathway is likely to function in parallel to an ATFS-1 dependent pathway. Individually, both chronic nicotine and *atfs-1* knockdown increase survival of DNs. However, their combined effects are deleterious. This suggests that the processes activated by both perturbations have a bell-shaped effect on survival. Indeed, two possible downstream targets affected by mitochondrial activity, cytosolic calcium and free radicals, also exhibit such bell-shaped effects on survival (Calì, Ottolini, & Brini, 2012). Both are required for survival at low concentrations and are toxic at higher concentrations.

Involvement of protective mitochondrial-dependent pathways in nicotine-mediated protection is consistent with recent analysis of a *CHRNA6* mutation. This mutation was identified as a genetic modifier that enhanced severity of infantile Parkinsonism and is likely to reduce activity of a DN-expressed nAChR. Importantly this mutation was also identified as an enhancer of the disease causing effects of a mutation in *WARS2*, mitochondrial tryptophanyl tRNA synthase, which was shown to reduce mitochondrial energy production (Martinelli et al., 2020).

Together, our results demonstrate that *C. elegans* represents a simple and well characterized model system that facilitates mechanistic understanding of nicotine-induced neuroprotection. Our results suggest that an interaction between nAChRs and D3Rs mediates a nicotine-dependent increase in cytosolic calcium. This, in turn, leads to calcium influx into mitochondria through the mitochondrial calcium uniporter, MCU-1, to produce mild mitochondrial stress, sufficient for activation of PINK-1-dependent protective processes within DNs (Figure 7).

Two findings from this research may explain selective protection of *substantia nigra* DNs by tobacco smoking. First, effects of nicotine require DOP-2, a D3R homolog, which is selectively expressed in DNs. Second, activation of PINK1-dependent mitochondrial quality control may selectively protect *substantia nigra* DNs; as these neurons were suggested to be uniquely susceptible to mitochondrial stress. Furthermore, many of the environmental and genetic risk factors for PD affect mitochondrial function (Ge et al., 2020).

In all, this research reveals a functional intersection of nicotinic receptor activity, dopaminergic neurotransmission and mitochondrial dynamics with a capacity to selectively modulate DN survival, and its clinical manifestations in PD. While tobacco smoking itself will certainly never be a panacea for PD, the molecular underpinnings of how it mysteriously confers protection provide for a unique and potentially valuable entreé into unexplored therapeutic strategies to combat degeneration of DNs.

## Materials and methods

### Strains

Nematodes were maintained using standard procedures (Brenner et al., 1974). The wild-type is Bristol N2. Additional *C. elegans* strains examined included nAChR subunit mutants: *unc-38(e264), unc-63(x37), acr-12(ok367), acr-8(ok1240), acr-2(ok1887), lev-1(e211), unc-29(e1072), acr-15(ok1214), acr-21(ok1314), acr-18(ok1285), deg-3des-2(u773)* (a deletion affecting both subunits), and *acr-5(ok180)*. All mutants examined in this study are considered *loss-of-function (lf)* mutants.

Analysis of DN survival following 6-OHDA exposure was examined using BY250 P_*dat-1*_::GFP (*vtIs7*), a kind gift from Randy Blakely (Florida Atlantic University). For the analysis of effects of nAChR mutations on DN survival, the mutations in the following nAChR subunit mutations were crossed into BY250: *unc-38(e264), unc-29(e1072), acr-2(ok1887) and lev-1(e211)*.

For DN-specific RNAi knockdown of genes, we used a *sid-1* mutant strain expressing *sid-1* only in DNs; this strain is UA202 [*sid-1(pk3321)*; P_*dat-1*_::GFP::*sid-1*, P_*myo-2*_::mCherry #15.1[*baIn36*]; P_*dat-1*_::GFP::GFP(*vtIs7*) (Harrington et al., 2012). RNAi clones were obtained from the Ahringer library (Kamath et al., 2003) except for *atfs-1*, which was described previously (Kamath et al., 2003; Martinez et al., 2017).

For mitochondrial stress assays, we used a chromosomally-integrated transcriptional reporter strain that expresses GFP under control of the *hsp-6* promoter (SJ4100 [*hsp-6::*GFP (*zcIs13*)]) (Yoneda et al., 2004).

### Basal slowing assay (BSR)

Basal slowing experiments were done as previously described (Chase et al., 2004; Sawin et al., 2000). In these experiments movement of animals on food is compared to their movement off food. Experiments were done on fresh NGM plates. For food containing plates we added 100μl of a ten-fold concentrated freshly grown late logarithmic stage HB101 bacteria. Bacteria were gently spread on the plates, air dried and then transferred to 37°C for overnight growth. Empty plates were also maintained overnight at 37°C. Animals for experiments were picked at the fourth larval stage to fresh plates for overnight growth at 20°C. Prior to the experiment plates were equilibrated at 20°C. Animals were transferred to plates with or without food using a glass capillary containing M9. Each animal was gently washed of bacteria before transfer to the new plate. 90 seconds after transfer to the experimental plate each animal was examined for appearance of body bends at the head. For each animal the number of sinusoidal bends initiating at the head and propagating backwards was counted for two consecutive 20 second periods, and the number given for each animal is the average of these two measurements. For experiments on food animals were placed in the middle of the bacterial lawn, and animals that moved to the edge of this lawn or that exited the food lawn were not analyzed.

### Exogenous dopamine response assay

Experiments were done as previously described (Chase et al., 2004). Dopamine containing plates were prepared the day prior to assay. These plates contained 100 ml double distilled water, 1.7gr agar, 11.4μl acetic acid and 0.379 gr dopamine hydrochloride (Abcam, ab1210565-5-B) to a final concentration of 20mM. Dopamine was added after the agar cooled down to 50°C. Prior to the experiment plates were air dried for one-hour. For experiments fourth larval stage animals were picked to fresh plates for overnight growth at 20°C. For analysis of the dopamine response 10 animals were placed on each dopamine plate and inspected for movement every 15 minutes for one-hour. Temperature for experiments was maintained at approximately 20°C. Animals were considered to be paralyzed if for 15 seconds no sinusoidal wave was seen in the head region. Animals showing only small head movements were considered paralyzed. In each experiment N2 controls were examined in parallel to mutant animals.

### RIC-3 rescue of the basal slowing response

For expression of *ric-3* in DNs only, we PCR amplified a 0.8 Kb fragment containing the *dat-1* promoter using the following primers (5’ to 3’): Dat1draIIIF, GATCACGTAGTGCTTCCATGAAATGG and Dat1sphIR, CAGCTTGCATGCACAAACTTGTATC. These primers include a *draIII* or a *SpHI* sites to enable cloning of the resulting PCR product upstream to a RIC-3::GFP fusion lacking the non-conserved coiled coil domain (Biala, Liewald, Cohen Ben-Ami, Gottschalk, & Treinin, 2009). The resulting plasmid was injected at 5ng/μl concentration into *ric-3(md1181)* animals together with a coelomocyte marker, CC::RFP at 50ng/μl and supplemented with empty bluescript vector as carrier 100 ng/μl. For the basal slowing experiments transgenic L4 animals expressing RFP in their coelomocytes were picked the day prior to the experiment for overnight growth at 20°C.

### 6-OHDA treatment and nicotine exposures

Animals were analyzed for degeneration of DNs as described previously (S. Cao, Gelwix, Caldwell, & Caldwell, 2005; Ray, Martinez, Berkowitz, Caldwell, & Caldwell, 2014). Briefly, animals were synchronized and grown at 20°C on NGM plates seeded with OP-50 bacteria. On day-3 post-hatching, animals were exposed to 10mM 6-OHDA/2mM ascorbic acid (Tocris Bioscience) for one-hour at 20°C. After the one-hour incubation, 6-OHDA was oxidized by adding 0.5X M9 and washed three times with ddH_2_O then plated back onto fresh, appropriate plates matching their respective conditions. Twenty-four hours after exposure, three replicates of 30 worms were scored for dopaminergic neurodegeneration of the six anterior DNs, totaling to 90 worms per treatment. Worms were scored as normal if no degenerative phenotypes (dendritic blebbing, swollen soma, broken projections, or a missing neuron) were observed in the six anterior [four CEPs (cephalic) and two ADEs (anterior deirid)] DNs.

Animals were treated with nicotine (Sigma N3876) using three different exposure paradigms – chronic, acute, and pre-exposure. These were performed as follows: 1) for chronic nicotine exposure, animals were raised on NGM plates with nicotine added to the agar to the final concentration of 62μM and nicotine was added to the 6-OHDA assay mix. Animals were placed back onto nicotine containing plates post-6-OHDA treatments; 2) for acute nicotine tests, animals were only exposed to nicotine while in the 6-OHDA assay mix; 3) for pre-exposure, a one-hour nicotine treatment was done 24-four hours prior to 6-OHDA, in addition to nicotine being present in the 6-OHDA assay mix. Animals in the control groups were never exposed to nicotine. The solvent used for nicotine was ddH_2_O.

### DN-specific knockdown of genes

RNAi knockdown of genes expressed in the DNs was accomplished as previously described (Harrington et al., 2012; Ray et al., 2014). Briefly, bacterial RNAi feeding constructs were obtained from the Ahringer *C. elegans* library (Kamath et al., 2003). Bacteria were isolated and grown overnight in Luria broth (LB) media containing 100μg/ml ampicillin. Nematode growth media (NGM) plates containing 50μg/ml ampicillin, 1mM IPTG, with or without 62μM nicotine, were seeded with 300μl of RNAi bacterial culture, allowed to dry and grown overnight. UA202 (DN-specific RNAi sensitive) animals were grown on RNAi conditions for two generations (Ray et al., 2014). The first generation was raised on RNAi plates without nicotine. Their embryos were collected and synchronized by hypochlorite treatment (Lewis and Fleming, 1995) and then placed onto agar plates corresponding to their respective RNAi conditions. Experimental plates had 62μM nicotine added to the agar for a chronic exposure (see above methods) while control treatments were kept off of nicotine for the entirety of the experiment. This method allows for stronger silencing of targeted genes without the nicotine-induced alteration of gene expression in the first generation. The second generation was exposed to 10mM 6-OHDA/2mM ascorbic acid at day-3 post-hatching for one-hour and then plated back onto RNAi-nicotine plates of their respective condition. They were then scored 24-hours later for DN neurodegeneration.

### Mitochondrial stress assay

Experiments testing the influence of nicotine on mitochondrial stress used the reporter strain SJ4100 (*hsp-6*::GFP (*zcIs13*)) (Yoneda et al., 2004). Animals were raised and kept on NGM plates seeded with OP-50 bacteria that had nicotine added to the agar to a final concentration of 62μM; control plates had no nicotine. GFP fluorescent images of 2-3 replicates, with 30 animals per replicate, were taken on days four and seven post-hatching. Focusing on an anatomically consistent region measuring 275 x 275 pixels comprising the same defined area in each animal (muscular region directly posterior to the pharynx), images were taken and pixel intensity was measured using MetaMorph software.

### Statistical Analysis

Data are represented as SD for all figures, “n” is number of animals examined and “N” is number of independent experiments or plates in dopamine experiments. Significance in BSR experiments were examined by two different methods: 1) one-way ANOVA with Tukey’s *post hoc* multiple comparisons test was used to compare the number of body bends; 2) unpaired two-tailed Student’s *t*-test was used to compare BSR index (number of bend bends on food relative to off food). Unpaired two-tailed Student’s *t*-test was used to compare paralyzing effects of exogenous dopamine. Pixel intensity comparisons used unpaired two-tailed Student’s *t*-test with 3 replicates of 30 worms per replicate (day seven used 2 replicates). All neurodegeneration experiments included 3 replicates of 30 worms per replicate and comparisons done by two-way ANOVA with Tukey’s *post hoc* multiple comparisons test. *p* < 0.05 was used the minimum threshold to determine statistical significance. Statistical comparisons were accomplished on GraphPad Prism.

## Acknowledgements

We would like to thank the members of the Caldwell and Treinin laboratories for their collegiality and helpful discussions surrounding this research. Special thanks to Dr. Laura Berkowitz for her assistance with experimental design and technical advice. JBN and GAC were supported by funds from NIH grant (R15NS104857-01A1), GH, AM, AK and MT were supported by a Prusiner-Abramsky research award and KAC was supported by NIH grant (R15NS106460-01A1).

